# The rate and molecular spectrum of spontaneous mutations in the GC-rich multi-chromosome genome of *Burkholderia cenocepacia*

**DOI:** 10.1101/011841

**Authors:** Marcus M. Dillon, Way Sung, Michael Lynch, Vaughn S. Cooper

## Abstract

Spontaneous mutations are ultimately essential for evolutionary change and are also the root cause of many diseases. However, until recently, both biological and technical barriers have prevented detailed analyses of mutation profiles, constraining our understanding of the mutation process to a few model organisms and leaving major gaps in our understanding of the role of genome content and structure on mutation. Here, we present a genome-wide view of the molecular mutation spectrum in *Burkholderia cenocepacia*, a clinically relevant pathogen with high %GC-content and multiple chromosomes. We find that *B. cenocepacia* has low genome-wide mutation rates with insertion-deletion mutations biased towards deletions, consistent with the idea that deletion pressure reduces prokaryotic genome sizes. Unlike prior studies of other organisms, mutations in *B. cenocepacia* are not AT-biased, which suggests that at least some genomes with high %GC-content experience unusual base-substitution mutation pressure. Importantly, we also observe variation in both the rates and spectra of mutations among chromosomes and elevated G:C>T:A transversions in late-replicating regions. Thus, although some patterns of mutation appear to be highly conserved across cellular life, others vary between species and even between chromosomes of the same species, potentially influencing the evolution of nucleotide composition and genome architecture.

## INTRODUCTION

As the ultimate source of genetic variation, mutation is implicit in every aspect of genetics and evolution. However, as a result of the genetic burden imposed by deleterious mutations, remarkably low mutation rates have evolved across all of life, making detection of these rare events technologically challenging and accurate measures of mutation rates and spectra exceedingly difficult (Kibota and Lynch 1996; Lynch and Walsh 1998; Sniegowski *et al.* 2000; Lynch 2011; Fijalkowska *et al.* 2012; Zhu *et al.* 2014). Until recently, most estimates of mutational properties have been derived indirectly using comparative genomics at putatively neutral sites (Graur and Li 2000; Wielgoss *et al.* 2011) or by extrapolation from small reporter-construct studies (Drake 1991). Both of these methods are subject to potentially significant biases, as many putatively neutral sites are subject to selection and mutation rates can vary substantially among different genomic regions (Lynch 2007).

To avoid the potential biases of these earlier methods, pairing classic mutation accumulation (MA) with whole-genome sequencing (WGS) has become the preferred method for obtaining direct measures of mutation rates and spectra (Lynch *et al.* 2008; Denver *et al.* 2009; Ossowski *et al.* 2010; Lee *et al.* 2012; Sung *et al.* 2012a; b; Heilbron *et al.* 2014). Using this strategy, a single clonal ancestor is used to initiate several replicate lineages that are subsequently passaged through repeated single-cell bottlenecks for several thousand generations. The complete genomes of each evolved lineage are then sequenced and compared with the other lines to identify *de novo* mutations occurring over the course of the experiment. The bottlenecking regime minimizes the ability of natural selection to eliminate deleterious mutations, and the parallel sequencing provides a large enough body of information to yield a nearly unbiased picture of the natural mutation spectrum of the study organism (Lynch *et al.* 2008).

The MA-WGS method has now been used to examine mutational processes in several model eukaryotic and prokaryotic species, yielding a number of apparently generalizable conclusions about mutation rates and spectra. For example, a negative scaling between base-substitution mutation rates and both effective population size (N_e_) and the amount of coding DNA supports the hypothesis that the refinement of replication fidelity that can be achieved by selection is determined by the power of random genetic drift among phylogenetic lineages (Lynch 2011; Sung *et al.* 2012a). This “drift-barrier hypothesis” therefore predicts that organisms with very large population sizes such as some bacteria should have evolved very low mutation rates (Lee *et al.* 2012; Sung *et al.* 2012a; Foster *et al.* 2013). Universal transition and G:C>A:T biases have also been observed in all MA studies to date (Lind and Andersson 2008; Lynch *et al.* 2008; Denver *et al.* 2009; Ossowski *et al.* 2010; Lee *et al.* 2012; Sung *et al.* 2012a; b), corroborating previous findings using indirect methods (Hershberg and Petrov 2010; Hildebrand *et al.* 2010). However, several additional characteristics of mutation spectra vary among species (Lynch *et al.* 2008; Denver *et al.* 2009; Ossowski *et al.* 2010; Lee *et al.* 2012; Sung *et al.* 2012a; b), and examining the role of genome architecture, size, and lifestyle in producing these idiosyncrasies will require a considerably larger number of detailed MA studies. Among bacterial species that have been subjected to mutational studies, genomes with high %GC-content are particularly sparse and no studies have been conducted on bacteria with multiple chromosomes, a genome architecture of many important bacterial species (e.g *Vibrio*, *Brucella*, *Burkholderia*).

*Burkholderia cenocepacia* is a member of the *Burkholderia cepacia* complex, a diverse group of bacteria with important clinical implications for patients with cystic fibrosis (CF), where they can form persistent lung infections and highly resistant biofilms (Coenye *et al.* 2004; Mahenthiralingam *et al.* 2005; Traverse *et al.* 2013). The core genome of *B. cenocepacia* HI2424 has a high %GC-content (66.8%) and harbors three chromosomes, each containing rDNA operons (LiPuma *et al.* 2002), although the third chromosome can be eliminated under certain conditions (Agnoli *et al.* 2012). The primary chromosome (Chr1) is ∼3.48 Mb and contains 3253 genes; the secondary chromosome (Chr2) is ∼3.00 Mb and contains 2709 genes; and the tertiary chromosome (Chr3) is ∼1.06 Mb and contains 929 genes. In addition, *B. cenocepacia* HI2424 contains a 0.164 Mb plasmid, which contains 159 genes and lower %GC-content than the core genome (62.0%). Although the %GC-content is consistent across the three core chromosomes, the proportion of coding DNA declines from Chr1 to Chr3, while the synonymous and non-synonymous substitution rates increase from Chr1 to Chr3 (Cooper *et al.* 2010; Morrow and Cooper 2012). Whether this variation in evolutionary rate is driven by variation in non-adaptive processes like mutation bias or variation in the relative strength of purifying selection remains a largely unanswered question in the evolution of bacteria with multiple chromosomes.

Here, we applied whole-genome sequencing to 47 MA lineages derived from *B. cenocepacia* HI2424 that were evolved in the near absence of natural selection for over 5550 generations each. We identified a total of 282 mutations spanning all three replicons and the plasmid, enabling a unique perspective on inter-chromosomal variation in both mutation rate and spectra, in a bacterium with the highest %GC-content studied with MA-WGS to date.

## MATERIALS AND METHODS

### Mutation accumulation

Seventy-five independent lineages were founded by single cells derived from a single colony of *Burkholderia cenocepacia* HI2424, a soil isolate that had only previously been passaged in the laboratory during isolation (Coenye and LiPuma 2003). Independent lineages were then serially propagated every 24 hours onto fresh high nutrient Tryptic Soy Agar (TSA) plates (30 g/L Tryptic Soy Broth (TSB) Powder, 15 g/L Agar). Two lineages were maintained on each plate at 37°C, and the isolated colony closest to the base of each plate half was chosen for daily restreaking. Following 217-days of MA, frozen stocks of all lineages were prepared by growing a final colony per isolate in 5 ml TSB (30 g/L TSB) overnight at 37°C, and freezing in 8% DMSO at -80°C.

Daily generation times were estimated each month by placing a single representative colony from each line in 2 ml of Phosphate Buffer Saline (80 g/L NaCl, 2 g/L KCl, 14.4 g/L Na_2_HPO_4_ • 2H_2_O, 2.4 g/L KH_2_PO_4_), serially diluting to 10^-3^ and spread plating 100 ul on TSA. By counting the colonies on the resultant TSA plate, we calculated the number of viable cells in a single colony and thus the number of generations between each transfer. The average generation time across all lines was then calculated and used as the daily generation time for that month. These generation-time measurements were used to evaluate potential effects of declining colony size over the course of the MA experiment as a result of mutational load, a phenotype that was observed (see Figure S1). Final generation numbers per line were estimated as the sum of monthly generation estimates, which were derived by multiplying the number of generations per day in that month by the number of days between measurements (see Figure S1).

### DNA extraction and sequencing

Genomic DNA was extracted from 1 ml of overnight culture inoculated from 47 frozen derivatives of MA lines and the ancestor of the MA experiments using the Wizard Genomic DNA Purification Kit (Promega Inc.). Following library preparation, sequencing was performed using the 151-bp paired-end Illumina HiSeq platform at the University of New Hampshire Hubbard Center for Genomic Studies with an average fragment size between paired-end reads of ∼386 bp. All of our raw fastQ files were analyzed using fastQC, but filtering was not performed as all files passed each of the fastQC modules. The average Phred score was 34 for both forward and reverse reads, with moderate declines in base quality from the beginning to the end of reads. All forward and reverse reads for each isolate and the ancestor were individually mapped to the reference genome of *Burkholderia cenocepacia* HI2424 (LiPuma *et al.* 2002), with both the Burrows-Wheeler Aligner (BWA) (Li and Durbin 2009) and Novoalign (<u>www.novocraft.com</u>), producing an average sequence depth of ∼50x.

### Base-substitution mutation identification

To identify base-substitution mutations, the sam alignment files that were produced by each reference aligner were first converted to mpileup format using samtools (Li *et al.* 2009). Forward and reverse read alignments were then produced for each position in each line using in-house perl scripts. Next, a three-step process was used to detect polymorphisms. First, an ancestral consensus base was called at each site in the reference genome as the base with the highest support among pooled reads across all lines, as long as there were at least three lines with sufficient coverage to identify a base. Importantly, this allows us to correct any differences between the published *B. cenocepacia* HI2424 reference and our ancestor, leveraging the enormous power of having sequenced 48 nearly isogenic isolates. Second, lineage specific consensus bases were called using the reads for each individually sequenced isolate, as long as the site was covered by at least two forward and two reverse reads, and at least 80% of those reads identified the same base. Lastly, these informative sites shared by MA lines and the ancestor were compared to identify putative base-substitution mutations. These mutations were considered genuine if both aligners independently identified the mutation and they were only identified in a single lineage. Base-substitutions shared by more than one lineage were considered to have occurred in the founding colony and were therefore only counted once, though this was only observed in two cases. Sites that did not meet the criteria for being informative in both an individual line and the ancestor were excluded from our analyses and do not contribute to the value of sites analyzed (*n*), described below for mutation rate calculations.

Although the above criteria for identifying individual line bases and overall consensus bases are relatively lenient given our coverage of ∼50x for individual lines, both the coverage and support for all substitutions that were called dramatically exceeded these criteria, demonstrating that we were not simply obtaining false positives in regions of lower coverage (see File S1). Furthermore, our ancestral strain was sequenced at the same depth as the derived strains and included as a 48^th^ isolate in our analysis. As expected, no mutations were identified in the ancestral strain. These same methods have been used to identify base-substitution mutations in both *Escherichia coli* and *Bacillus subtilis* MA lines, where 19 of 19 and 69 of 69 base-substitution mutations called were confirmed by conventional sequencing, respectively (Lee *et al.* 2012; Sung *et al.* 2012a). Thus, these criteria are unlikely to result in false positives, while allowing us to cover the majority of the *B. cenocepacia* genome and reduce false negatives.

### Insertion-deletion mutation identification

For insertion-deletion mutations (indels), inherent difficulties with gaps and repeat elements can reduce agreement in the alignment of single reads using short-read alignment algorithms, even in the case of true indels. Thus, putative indels were first extracted from both BWA and Novoalign at all sites where at least two forward and two reverse reads covered an indel, and 30% of those reads identified the exact same indel (size and motif). Next, the alignment output was additionally passaged through the pattern-growth algorithm PINDEL to verify putative indels from the alignment and identify larger indels using paired-end information (Ye *et al.* 2009). Here, a total of twenty reads, including at least six forward and six reverse reads were required to extract a putative indel. Putative indels were only kept as true indels for further analysis if: a) they were independently identified by both alignment algorithms and PINDEL, and at least 50% of the full-coverage reads (>25 bases on both sides of the indel) from the initial alignment identified the mutation; b) they were identified only by BWA and Novoalign, and at least 80% of the good-coverage reads from the initial alignment identified the mutation; or c) they were larger indels that were only identified by the more strict requirements of PINDEL. Moreover, if an indel was identified in more than half of the lineages, we consider it to be an ancestral indel and exclude it from further analyses.

Unlike base-substitutions mutations, many reads that cover an indel mutation may fail to identify the mutation because they lack sufficient coverage on both sides of the mutation to anchor the read to the reference genome, particular when they occur in simple sequence repeats. Therefore, applying the initially lenient filter to extract putative indels is justified to identify all potential indels. By then focusing only on the good-coverage reads and applying an independent paired-end indel identifier (PINDEL), we can filter out indels that are more likely to be false positives, while keeping only the high concordance indels supported by multiple algorithms. Although there remains more uncertainty with indel calls than with base-substitutions mutations, we are confident that we have obtained an accurate picture of the naturally occurring indels from this study because of the high concordance across algorithms and reads (see File S1; Figure S3), and the fact that no indels were called independently by more than two lines (see Figure S4). A complete list of the indels identified in this study, along with the algorithms that identified them, their coverage, and concordance across well-covered reads can be found in File S1.

### Mutation-rate analysis

Once a complete set of mutations had been identified in each lineage, we calculated the substitution and indel mutation rates for each line using the equation μ = *m*/ *nT*, where *μ* represents the mutation rate (*μ*_*bs*_ for bps, *μ*_*indel*_ for indels), m represents the number of mutations observed, n represents the number of sites that had sufficient depth and consensus to analyze, and T represents the total generations over the course of the MA study for an individual line. The standard error of the mutation rate for each line was measured as described previously with the equation 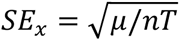 (Denver *et al.* 2004, 2009).

The final *μ*_*bs*_ and *μ*_*indel*_ for *B. cenocepacia* were calculated by taking the average *μ* of all sequenced lineages, and the total standard error was calculated as the standard deviation of the mutation rates across all lines (*s*) divided by the square root of the number of lines analyzed 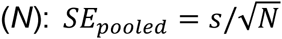. Specific base-substitution mutation rates were further divided into conditional rates for each substitution type, again using the equation *μ*_*bs*_= *m*/*nT,* where *m* is the number of substitutions of a particular type, and *n* is the number of ancestral bases that can lead to each substitution with sufficient depth and consensus to analyze. The conditional substitution rates at seven MLST loci (*atpD*, *gltB*, *gyrB*, *lepA*, *phaC*, *recA*, and *trpB*) were calculated under the assumption that the most common nucleotide was the ancestral state and any deviation from that ancestral state occurred only once and spread through the population (Jolley and Maiden 2010). We then estimated conditional substitution rates as *μ*_*b,s*_ = *m*/*n,* as described above.

### Calculation of *G*_*E*_, *π*_*s*_, and *N*_*E*_

Effective genome size (*G*_*E*_) was determined as the total coding bases in the *B. cenocepacia* genome. Silent site diversity (*π*_*s*_) was derived using the MLST loci described above, which were concatenated and aligned using BIGSdb (Jolley and Maiden 2010), and analyzed using DNAsp (Librado and Rozas 2009). Using the value of *μ*_*bs*_ obtained in this study, N_e_ was estimated by dividing the value of *π*_s_ by 2*μ*_*bs*_ (π_*s*_ = 2*N*_*e*_μ_*bs*_) (Kimura 1983).

## RESULTS

A classic mutation-accumulation experiment was carried out for 217 days with 75 independent lineages all derived from the same ancestral colony of *B. cenocepacia* HI2424 (LiPuma *et al.* 2002). This method thus founds a new population each day by a single cell, which limits the efficiency with which natural selection can purge deleterious and enrich beneficial mutations. Measurements of generations of growth per day were taken monthly and varied from 26.2 ± 0.12 to 24.9 ± 0.14 (mean ± 95% CI of highest and lowest measurements, respectively) (see Figure S1), resulting in an average of 5554 generations per line over the course of the MA experiment. Thus, across the 47 lines whose complete genomes were sequenced, we were able to visualize the natural mutation spectrum of *B. cenocepacia* HI2424 over 261,047 generations of MA.

From the comparative sequence data, we identified 245 base-substitutional (bps) changes, 33 short-insertion/deletion (indel) mutations (with sizes in the range of 1 to 145 base pairs), and four plasmid-loss events spanning the entire genome (Figure 1). With means of 5.21 bps and 0.70 indel mutations per line, the distribution of bps and indels across individual lines did not differ significantly from a Poisson distribution (bps: χ^2^ = 1.81, p = 0.99; indels: χ^2^ = 0.48, p = 0.92), indicating that mutation rates did not vary over the course of the MA experiment (see Figure S2).

**Figure 1.**
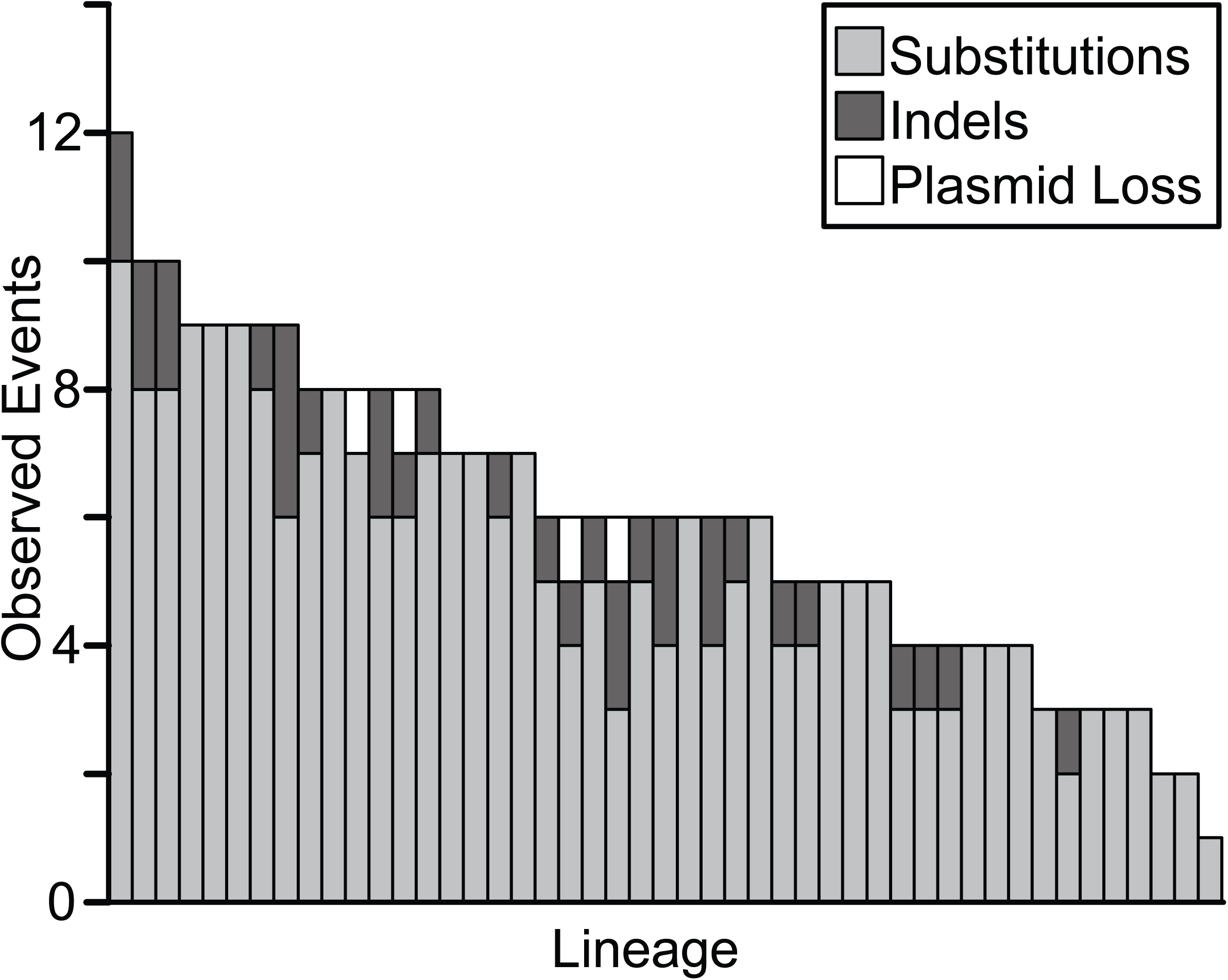
Distribution of observed mutations in the 47 sequenced lineages derived from *B. cenocepacia* HI2424 following an average of 5554 generations of mutation accumulation per line. Minimal variation exists in the number of sites analyzed per line, making differences in mutation number across lineages representative of the random variance in mutation rate measurements across lineages.

Mutation-accumulation experiments rely on the basic principle that when the effective population size (N_e_) is sufficiently reduced, the efficiency of selection is minimized to the point at which nearly all mutations become fixed by genetic drift with equal probability (Kibota and Lynch 1996). N_e_ in this mutation accumulation study was calculated to be ∼12.86, using the harmonic mean of the population size over 24 hours of colony growth (Hall *et al.* 2008). The threshold selective coefficient below which genetic drift will overpower natural selection is N_e_ × s = 1 for haploid organisms (Lynch 2007). Thus, only mutations conferring adaptive or deleterious effects of s > 0.078 would be subject to the biases of natural selection in this study, which is expected to be a very small fraction of mutations (Kimura 1983; Elena *et al.* 1998; Zeyl and DeVisser 2001; Hall *et al.* 2008).

Given the codon usage and %GC-content of synonymous and non-synonymous sites in *B. cenocepacia* HI2424, 27.8% of coding substitutions are expected to be synonymous in the absence of natural selection. The observed percentage of synonymous substitutions (25.5%) did not differ significantly from this null-expectation (χ^2^ = 0.54, df = 1, p = 0.46). Further, we find limited evidence of positive selection since parallel evolution among base-substitution mutations is rare in this study; no gene is hit more than twice across any of the 47 independently derived lineages (see File S1). Although both base-substitutions (χ^2^ = 4.20, df = 1, p = 0.04) and indels (χ^2^ = 21.3, df = 1, p < 0.0001) were biased to non-coding DNA, evidence exists that mismatch repair preferentially repairs damage in coding regions, which can create artificial signatures of selection in MA experiments (Lee *et al.* 2012). Thus, our overall observations are consistent with our MA experiment inducing limited selection on the mutation spectra; at least as far as base-substitutions are concerned.

### Low base-substitution and indel mutation rates

The preceding results imply that base-substitution and indel mutation rates for *B. cenocepacia* are 1.33 (0.008) × 10^-10^ /bp/generation and 1.68 (0.003) × 10^-11^ /bp/generation (SEM), respectively. Based on the 7.70 Mb genome size, these per-base mutation rates correspond to a genome-wide base-substitution mutation rate of only 0.001/genome/generation, and an indel mutation rate of only 0.0001/genome/generation. Although the ∼1:3 ratio of synonymous to non-synonymous substitutions is consistent with negligible influence of selection on base-substitution mutations in this study, too few indels occurred to evaluate a signature of selection, but their scarcity could reflect some selective loss of genotypes with loss-of-function mutations (Heilbron *et al.* 2014; Zhu *et al.* 2014).

### Base-substitution mutations are not AT-biased

One of the central motivations for studying the molecular mutation spectrum of *B. cenocepacia* was its high %GC-content (66.8%). A universal mutation bias in the direction of AT has been observed in all other wild-type species studied by MA (Table 1), and has also been inferred in comparative analyses of several bacterial species, including *Burkholderia pseudomallei* (Lynch *et al.* 2008; Denver *et al.* 2009, 2012; Keightley *et al.* 2009; Hershberg and Petrov 2010; Hildebrand *et al.* 2010; Lynch 2010a; Ossowski *et al.* 2010; Sung *et al.* 2012a; b; Lee *et al.* 2012; Schrider *et al.* 2013; Zhu *et al.* 2014). Thus, biased gene conversion and selection have been invoked to explain the high %GC-content realized in many genomes (Lynch *et al.* 2008; Duret and Galtier 2009; Raghavan *et al.* 2012; Zhu *et al.* 2014; Lassalle *et al.* 2015). Our data for *B. cenocepacia* are inconsistent with prior published studies showing a mutation bias in the direction of AT (Table 1), but also suggest that biased gene conversion and/or selection must have mostly generated the realized %GC-content of *B. cenocepacia*, which is substantially higher than expected based on mutation pressure alone.

**Table 1.**
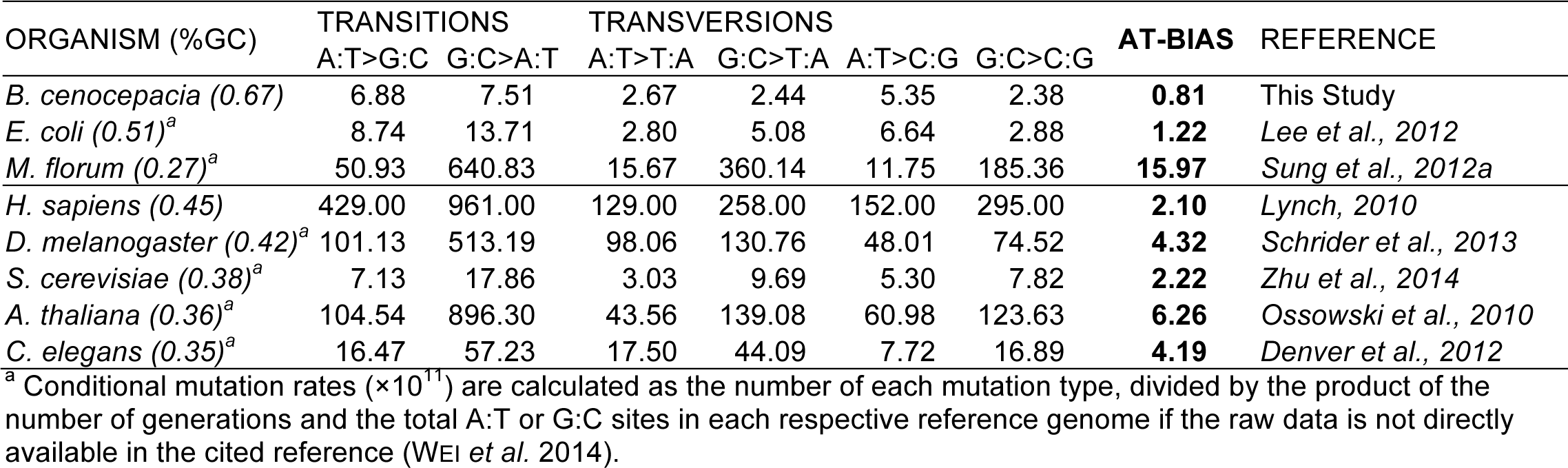
Comparison of mutation bias in *B. cenocepacia* with the mutation bias of seven other species with large mutation data sets and intact mismatch repair (two prokaryotes, five eukaryotes). The strength of the AT-mutation bias, calculated as the ratio of the conditional mutation rates in the in the G:C>A:T direction to the conditional mutation rates in the A:T>G:C direction, is substantially higher in all other species than in *B. cenocepacia*.

| ORGANISM (%GC) | TRANSITIONS |  | TRANSVERSIONS |  |  |  | AT-BIAS | REFERENCE |
| --- | --- | --- | --- | --- | --- | --- | --- | --- |
|  | A:T>G:C | G:C>A:T | A:T>T:A | G:C>T:A | A:T>C:G | G:C>C:G |  |  |
| *B. cenocepacia* (0.67) | 6.88 | 7.51 | 2.67 | 2.44 | 5.35 | 2.38 | **0.81** | This Study |
| *E. coli* (0.51)<sup>a</sup> | 8.74 | 13.71 | 2.80 | 5.08 | 6.64 | 2.88 | **1.22** | *Lee et al., 2012* |
| *M. florum* (0.27)<sup>a</sup> | 50.93 | 640.83 | 15.67 | 360.14 | 11.75 | 185.36 | **15.97** | *Sung et al., 2012a* |
| *H. sapiens* (0.45) | 429.00 | 961.00 | 129.00 | 258.00 | 152.00 | 295.00 | **2.10** | *Lynch, 2010* |
| *D. melanogaster* (0.42)<sup>a</sup> | 101.13 | 513.19 | 98.06 | 130.76 | 48.01 | 74.52 | **4.32** | *Schrider et al., 2013* |
| *S. cerevisiae* (0.38)<sup>a</sup> | 7.13 | 17.86 | 3.03 | 9.69 | 5.30 | 7.82 | **2.22** | *Zhu et al., 2014* |
| *A. thaliana* (0.36)<sup>a</sup> | 104.54 | 896.30 | 43.56 | 139.08 | 60.98 | 123.63 | **6.26** | *Ossowski et al., 2010* |
| *C. elegans* (0.35)<sup>a</sup> | 16.47 | 57.23 | 17.50 | 44.09 | 7.72 | 16.89 | **4.19** | *Denver et al., 2012* |

<sup>a</sup> Conditional mutation rates ( $\times 10^{11}$ ) are calculated as the number of each mutation type, divided by the product of the number of generations and the total A:T or G:C sites in each respective reference genome if the raw data is not directly available in the cited reference (Wei et al. 2014).

In comparing the relative rates of G:C>A:T transition and G:C>T:A transversion mutations with those of A:T>G:C transitions and A:T>C:G transversions, corrected for the ratio of G:C to A:T sites analyzed in this study, we found that substitutions in the G:C direction were 17% more frequent than mutations in the A:T direction per base pair, although the conditional rates were not significantly different (χ^2^ = 0.91, df = 1, p = 0.33). The lack of mutational bias in the A:T direction can largely be attributed to A:T>C:G transversions occurring at significantly higher rates than any other transversion type, most notably the G:C>T:A transversions (χ^2^ = 8.68, df = 1, p = 0.0032). However, A:T>G:C transitions also occurred at nearly the same rate as G:C>A:T transitions, the latter of which have been the most commonly observed substitution in other studies, putatively due to deamination of cytosine or 5-methyl-cytosine (Figure 2) (Lee *et al.* 2012; Sung *et al.* 2012b; Zhu *et al.* 2014).

**Figure 2.**
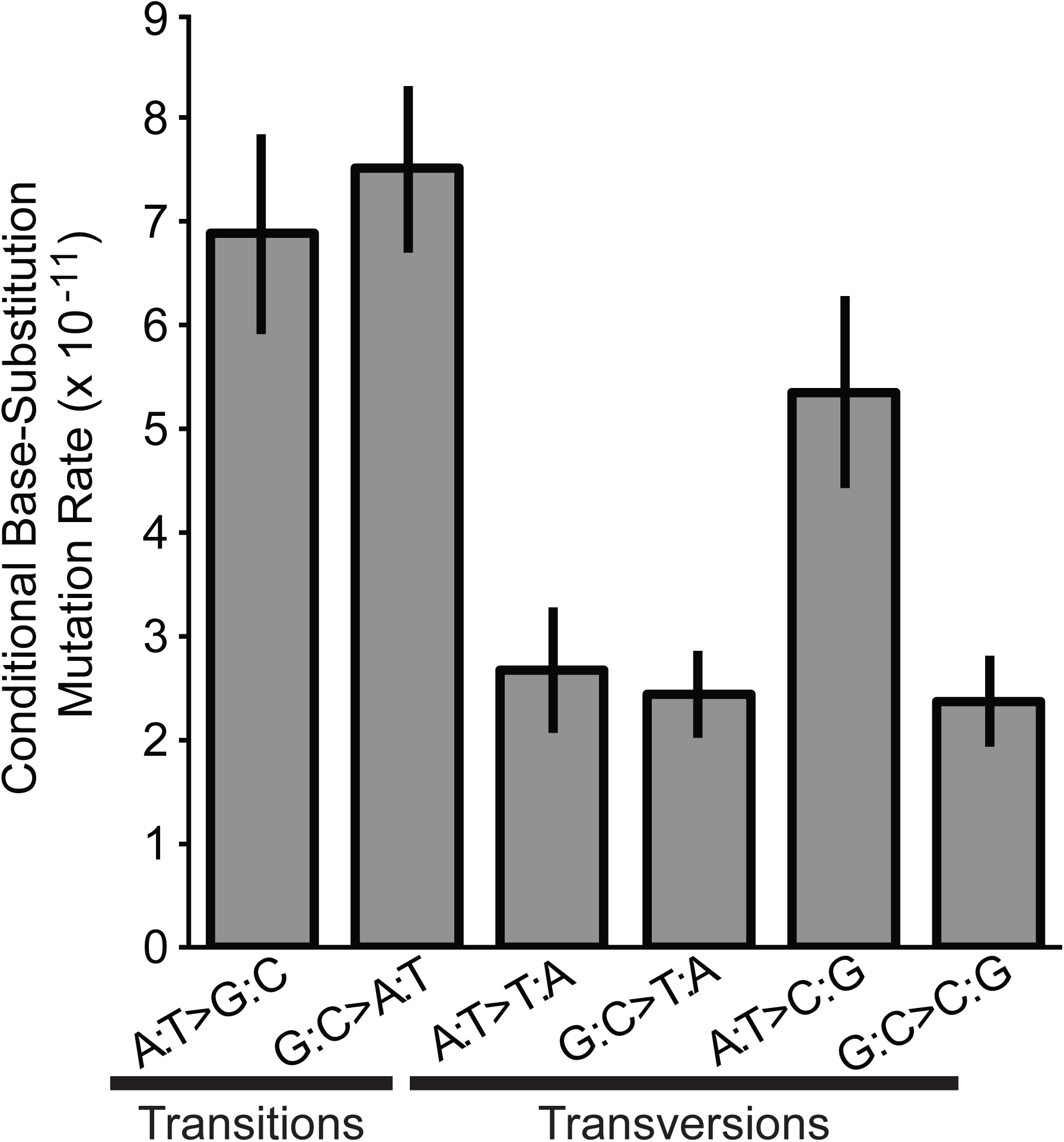
Conditional base-substitution mutation rates of *B. cenocepacia* mutation accumulation (MA) lines across all three chromosomes. Conditional base-substitution rates are estimated by scaling the base-substitution mutation rates to the analyzed nucleotide content of the *B. cenocepacia* genome, whereby only covered sites capable of producing a given substitution are used in the denominator of each calculation. Error bars represent one standard error of the mean.

Using the ratio of the conditional rate of mutation in the G:C direction to that in the A:T direction (x), the expected %GC-content under mutation-drift equilibrium is x/(1+x) = 0.539 ± 0.043 (SEM). Therefore, although mutation pressure in *B. cenocepacia* does not favor AT-bases, it is clear that the observed mutation bias is not sufficient to elicit the realized %GC-content of 66.8%. Thus, either the *B. cenocepacia* genome is still moving towards mutation-drift equilibrium, or GC-biased gene conversion and/or natural selection are responsible for the observed %GC-content (Lynch *et al.* 2008; Duret and Galtier 2009; Raghavan *et al.* 2012; Zhu *et al.* 2014; Lassalle *et al.* 2015).

### Deletion bias favors genome-size reduction and AT composition

Although our lower bound estimates of the insertion and deletion mutation rates are both ∼15-fold lower than the base-substitution mutation rate, many indels affect more than one base. Specifically, the 17 deletions observed in this study result in the deletion of a total of 376 bases, while the 16 insertions result in a gain of 121 bases. Therefore, the number of bases that are impacted by indels in this study is more than twice the number impacted by bps, indicating that indels may still play a central role in the genome evolution of *B. cenocepacia* if they are not purged by natural selection.

Although the ratio of deletions to insertions observed in this study was nearly 1, the per base-pair deletion rate (1.97 (0.86) × 10^-10^/bp/generation) was substantially higher than the insertion rate (6.11 (1.90) × 10^-11^/bp/generation), since the average size of deletions was greater than the average size of insertions. Thus, there is a net deletion rate of 1.36 (5.95) × 10^-10^/bp/generation (Table 2). Although no indels >150 bp were observed in this study, examining the depth of coverage of the *B. cenocepacia* HI2424 plasmid relative to the rest of the genome revealed that the plasmid was lost at a rate of 1.53 × 10^-5^ per cell division, while gains in plasmid copy number were not observed (Table 2).

**Table 2.** Parameters of insertion and deletion mutations during 261,038 generations of spontaneous mutation accumulation in *B. cenocepacia*.

| PARAMETER | DELETIONS | INSERTIONS |
| --- | --- | --- |
| Events Observed | 17 | 16 |
| Total Nucleotides Affected | 376 | 121 |
| Total GC Bases Affected | 302 | 87 |
| Total AT Bases Affected | 74 | 34 |
| Proportion of GC Bases Affected | 0.80 | 0.72 |
| Plasmid Copy Number Loss/Gain | 4 | 0 |

The base composition of deletions was also biased, with GC bases being deleted significantly more than expected based on the genome content (χ^2^ = 30.4, df = 1, p < 0.0001). In contrast, no detectable bias was observed towards insertions of GC over AT bases (χ^2^ = 1.20, df = 1, p = 0.27) (Table 2). Thus, indels in *B. cenocepacia* are expected to reduce genome wide %GC-content, further supporting the need for other population-genetic processes to account for the composition of high-GC genomes (Lynch *et al.* 2008; Duret and Galtier 2009; Raghavan *et al.* 2012; Zhu *et al.* 2014; Lassalle *et al.* 2015). Overall, the observed mutation spectra in this study suggest that the natural indel spectrum of *B. cenocepacia* causes both genome-size reduction and increased %AT-content.

### Non-uniform chromosomal distribution of mutations

Another major goal of this study was to investigate whether mutation rates and spectra vary among chromosomes and chromosomal regions. The three core chromosomes of *B. cenocepacia* vary in size and content but are sufficiently large to have each accumulated a considerable number of mutations in this study (Morrow and Cooper 2012). Chromosome 1 (chr1) is the largest chromosome (both in size and in gene count), with more essential and highly expressed genes than either chromosome 2 (chr2) or 3 (chr3) (see Figure S5). Expression and number of essential genes are second highest on chr2 and lowest on chr3 (Cooper *et al.* 2010; Morrow and Cooper 2012). In contrast, average non-synonymous and synonymous variation among orthologs shared by multiple strains of *B. cenocepacia,* as well as fixed variation among *Burkholderia* species (dN and dS), are highest on chr3 and lowest on chr1 (see Figure S5) (Cooper *et al.* 2010; Morrow and Cooper 2012).

The overall base-substitution mutation rates of the three core chromosomes differ significantly based on a chi-square proportions test, where the null expectation was that the number of substitutions would be proportional to the number of sites covered on each chromosome (χ^2^ = 6.77, df = 2, p = 0.034) (Figure 3A). Specifically, base-substitution mutation rates are highest on chr1, and lowest on chr2, which is the opposite of observed evolutionary rates on these chromosomes (see Figure S5) (Cooper *et al.* 2010). In addition, a second chi-squared test was performed to test whether the observed base-substitution mutation rates differed from the conditional mutation rates expected on each chromosome given their respective nucleotide contents, which are similar (%GC: Chr1-66.8%; Chr2-66.9%; Chr3-67.3%). Here, the null expectation for the total number of base-substitution mutations on each chromosome was calculated as the product of the number of GC bases covered, the total number of generations across lines, and the overall GC substitution rate across the genome, added to the product of the same calculation for AT substitutions. The differences in the base-substitution mutation rates of the three core chromosomes remained significant when this test was performed (χ^2^ = 6.88, df = 2, p = 0.032), indicating that the intrachromosomal heterogeneity in base-substitution mutation rates cannot be explained by variation in nucleotide content.

**Figure 3.**
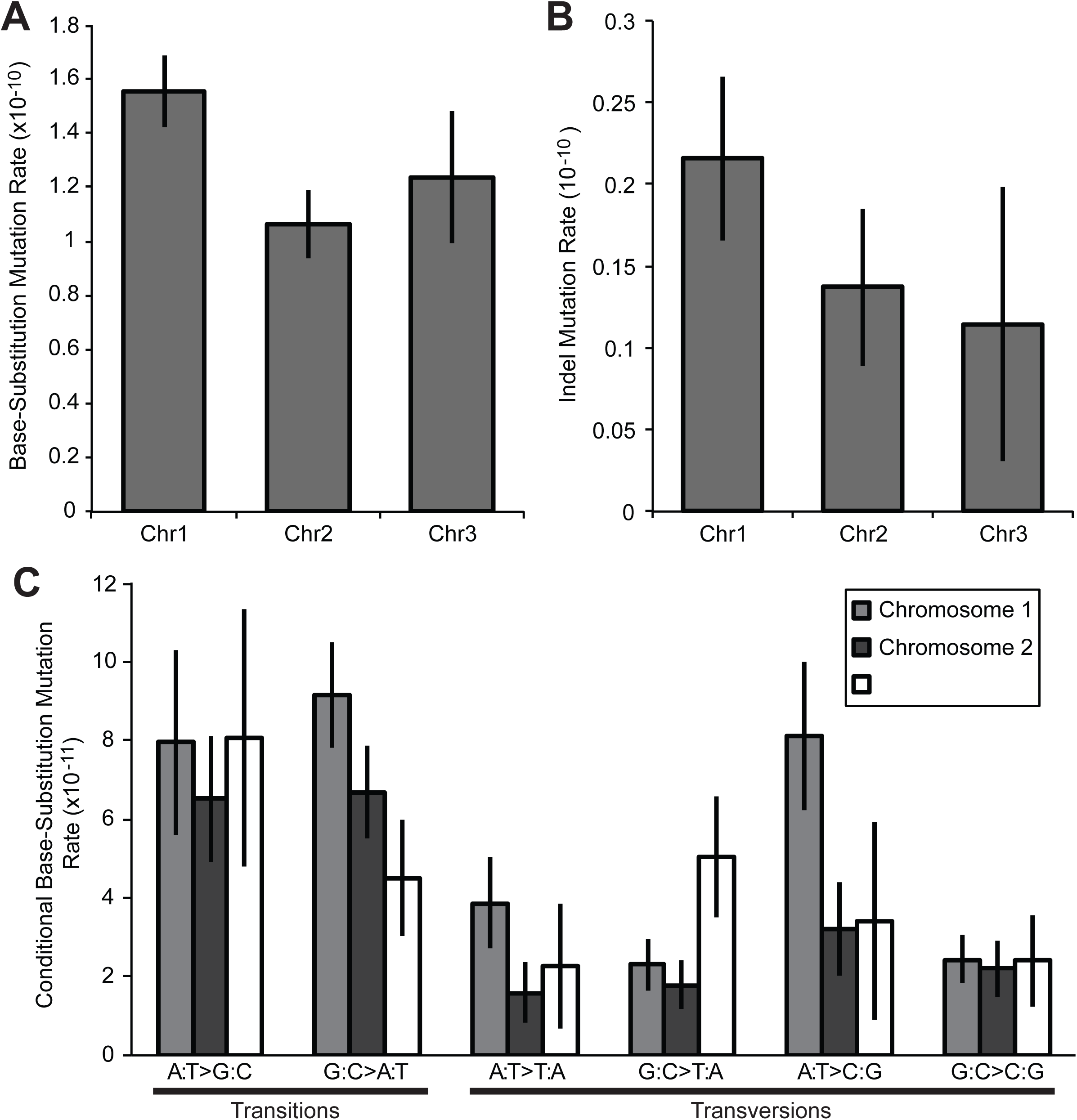
Substitution and indel mutation rates for the three chromosomes of *B. cenocepacia*; error bars indicate one standard error of the mean. A, B) Overall base-substitution and indel mutation rates. C) Conditional base-substitution mutation rates for each chromosome of *B. cenocepacia* estimated as described in Figure 2, based on the analyzed nucleotide content of each chromosome.

The conditional base-substitution mutation spectra were also significantly different in all pairwise chi-squared proportions tests between chromosomes (chr1/chr2: χ^2^=14.3, df=5, p=0.014; chr1/chr3: χ^2^=17.0, df=5, p=0.004; chr2/chr3: χ^2^=13.4, df=5, p=0.020) (Figure 3C). These comparisons further illustrate that the significant variation in conditional base-substitution mutation rates is mostly driven by a few types of substitutions that occur at higher rates on particular chromosomes. Specifically, although their individual differences were not quite statistically significant, G:C>T:A transversions seem to occur at the highest rate on chr3 (χ^2^ = 5.94, df = 2, p = 0.051) and A:T>C:G transversions occur at the highest rate on chr1 (χ^2^ = 5.67, df = 2, p = 0.059) (Figure 3B; Figure 4A).

**Figure 4.**
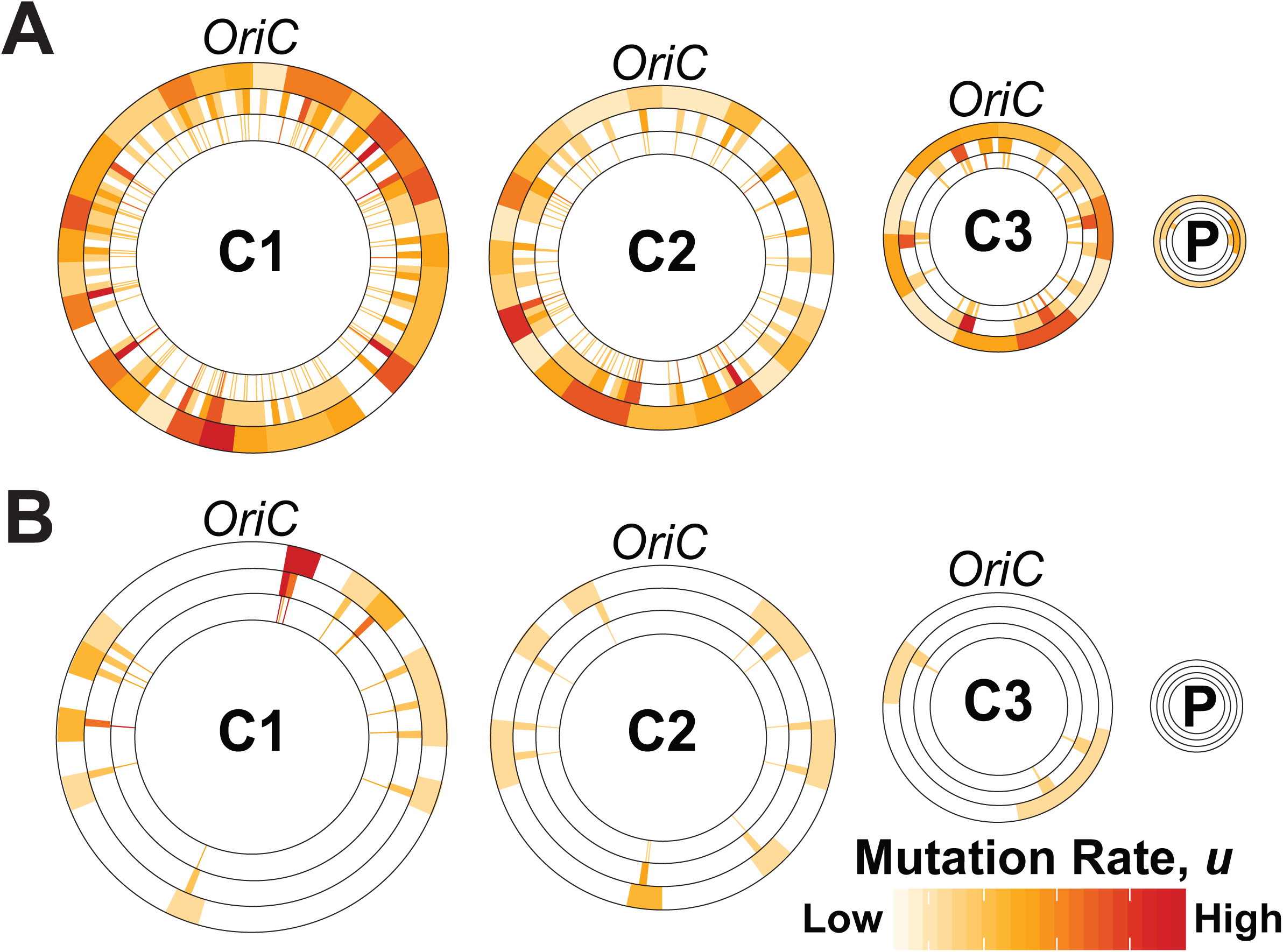
Overall base-substitution (A) and indel mutation rates (B) separated into 100 Kb (outer), 25 Kb (middle), and 5 Kb (inner) bins extending clockwise from the origin of replication (*oriC*) to reveal broad and local properties of intragenic variation in mutation rates of *B. cenocepacia*. Mutation rates were analyzed independently for each bin size, so color shades in smaller bins don’t directly compare to the same color shades in larger bins. The 0.164 Mb plasmid is not to scale.

Studies in *Vibrio cholerae* have suggested that in bacteria with multiple chromosomes, smaller secondary chromosomes delay their replication until there remains approximately the same number of bases to be replicated on larger chromosomes (Rasmussen *et al.* 2007; Cooper *et al.* 2010). This ensures synchrony of replication termination between chromosomes of different sizes, despite the fact that their replication proceeds at the same rate. To test whether this replication timing gradient is partially responsible for the patterns we observe in base-substitution mutation spectra between chromosomes, we binned chr1 and chr2 into late and early replicating regions, where the early replicating regions represent bases presumed to replicate prior to chr3 initiation, and the late replicated regions represent bases presumed to replicate following chr3 initiation (the last 1.06 Mb replicated).

In support of this model, G:C>T:A transversions also occur at a slightly higher rate in late replicated regions of chr1 and chr2 than they do in early replicated regions of chr1 and chr2 (see Figure 5A). However, even when mutations are binned by overall replication timing (combining late replicating regions on chr1 and chr2 with chr3 and comparing them to early replicating regions on chr1 and chr2), the rate of G:C>T:A transversions is not significantly higher than it is in early replication-timing regions, likely due to small sample sizes (χ^2^ = 2.52, df = 1, p = 0.113). A:T>C:G transversions occur at slightly higher rates in early replicated regions of chr1 and chr2 than they do in late replicated regions (see Figure 5B), but again the difference is not statistically significant (χ^2^ = 1.26, df = 1, p = 0.262). Together, these findings suggest that late replicating DNA is predisposed to incur more G:C>T:A transversions and early replicating DNA is predisposed to incur more A:T>C:G transversions, but a larger collection of mutations will be necessary to fully address this question.

**Figure 5.**
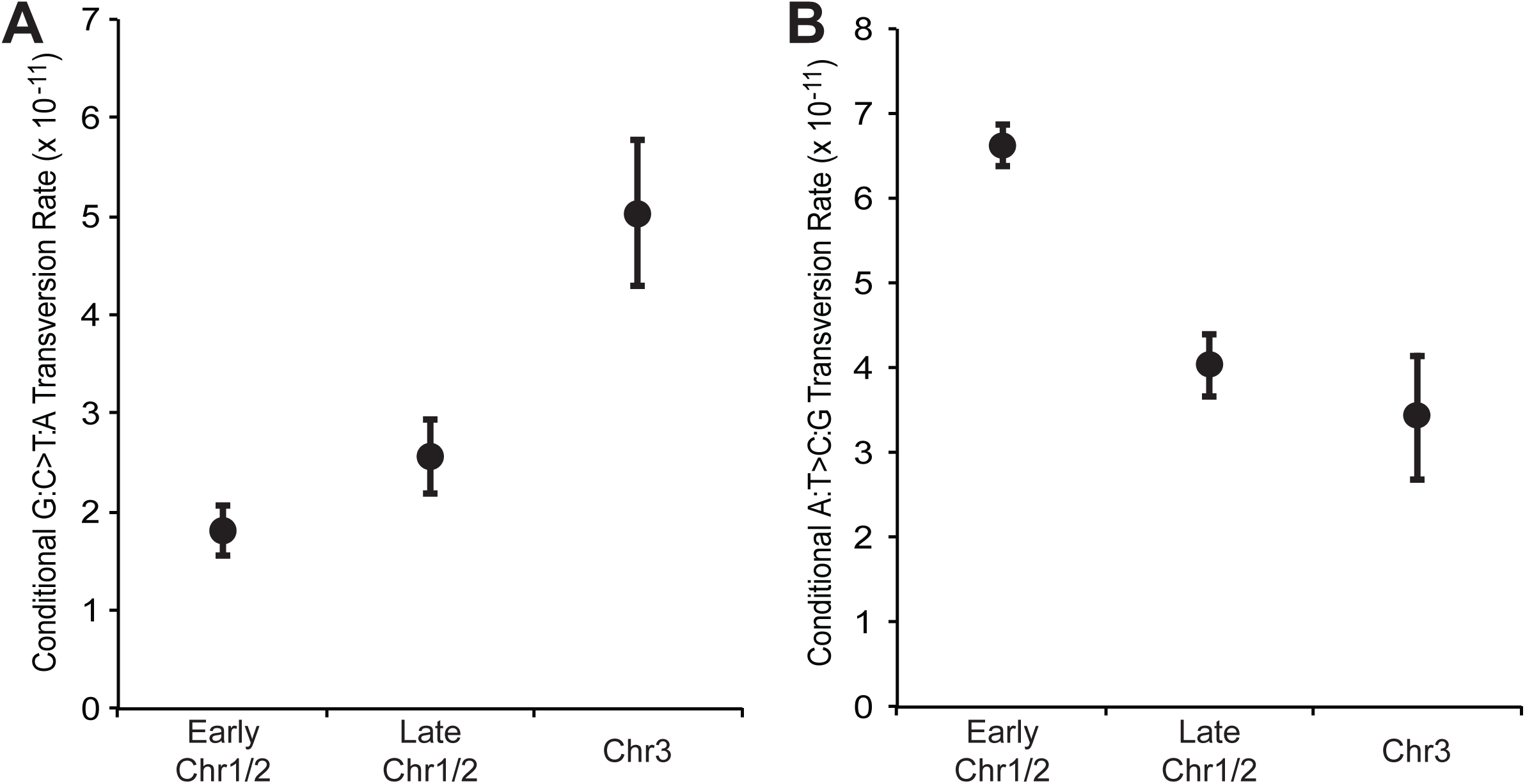
Conditional G:C>T:A (A) and A:T>C:G (B) transversion rates, normalized for base composition, between regions on the primary and secondary chromosomes that are replicated prior to initiation of replication of the third chromosome (Early Chr1/2), regions on the primary and secondary chromosomes that are replicated simultaneously with the third chromosome (Late Chr1/2), and the third chromosome itself (Chr3), based on models from (Rasmussen *et al.* 2007). Error bars indicate one SEM.

Unlike base-substitution mutation rates, neither the deletion or insertion mutation rate varied significantly among chromosomes (Deletions: χ^2^=3.81, df=2, p=0.15; Insertions: χ^2^=0.64, df=2, p=0.73), (Figure 3B; Figure 4B). No indels were observed on the 0.16 Mb plasmid, but as noted above, four plasmid loss events were observed. The latter events involve the loss of 157 genes, and are expected to have phenotypic consequences. The relative rarity of indels observed in this study limits our ability to analyze their intra-chromosomal biases in great detail, but the repeated occurrence of indels in short 5kb regions, and particularly within microsatellites (57.6% of all indels) suggests that replication slippage is a common cause of indels in the *B. cenocepacia* genome (Figure 4B).

## DISCUSSION

Despite their relevance to both evolutionary theory and human health, the extent to which generalizations about mutation rates and spectra are conserved across organisms remains unclear. Bacterial genomes are particularly amenable to studying these issues because of their diverse genome content (Lynch 2007). In measuring the rate and molecular spectrum of mutations in the high-GC, multi-replicon genome of *B. cenocepacia*, we have corroborated some prior findings of MA studies in model organisms, but also demonstrated idiosyncrasies in the *B. cenocepacia* spectrum that may extend to other organisms with high %GC-content and/or with multiple chromosomes. Specifically, *B. cenocepacia* has a relatively low mutation rate and a mutation spectrum consistent with a universal deletion bias in prokaryotes (Mira *et al.* 2001). However, the lack of AT-mutation bias is inconsistent with all previous findings in mismatch-repair proficient organisms (Lynch *et al.* 2008; Denver *et al.* 2009; Hershberg and Petrov 2010; Hildebrand *et al.* 2010; Ossowski *et al.* 2010; Lee *et al.* 2012; Sung *et al.* 2012b). Further, both mutation rates and spectra differed significantly among chromosomes in a manner suggesting greater oxidative damage or more inefficient repair in late replicated regions.

As a member of a species complex with broad ecological and clinical significance, *B. cenocepacia* is a taxon with rich genomic resources that enable comparisons between the *de novo* mutations reported here and extant sequence diversity. With 7050 genes, *B. cenocepacia* HI2424 has a large amount of coding DNA (G_E_) (6.8×10^6^ base pairs), and a high average nucleotide heterozygosity at silent-sites (π_s_) (6.57×10^-2^) relative to other strains (Watterson 1975; Mahenthiralingam *et al.* 2005). By combining this *π*_s_ measurement and the base-substitution rate from this study, we estimate that the N_e_ of *B. cenocepacia* is approximately 2.47×10^8^, which is in the upper echelon among species whose N_e_ has been derived in this manner (see Figure S6). Under the drift-barrier hypothesis, high target size for functional DNA and high N_e_ increase the ability of natural selection to reduce mutation rates (Lynch 2010b, 2011; Sung *et al.* 2012a). Thus, given the large proteome and N_e_ of *B. cenocepacia*, it is unsurprising that *B. cenocepacia* has relatively low base-substitution and indel mutation rates when compared to other organisms (Sung *et al.* 2012a). However, the low substitution and indel mutation rates observed in this study need not imply limited genetic diversity among species of the *Burkholderia cepacia* complex. Rather, because of their high N_e_ and evidently frequent lateral genetic transfer, species of the *Burkholderia cepacia* complex are remarkably diverse (Baldwin *et al.* 2005; Pearson *et al.* 2009), demonstrating that low mutation rates need not imply low levels of genetic diversity.

*Burkholderia* genomes also tend to be large in comparison to other Proteobacteria, but this is evidently not the product of more frequent insertions. Rather, insertions and deletions occurred at similar rates but deletions were larger than insertions, and plasmids were lost relatively frequently, which together add to the general model that bacterial genomes are subject to a deletion bias (Mira *et al.* 2001; Kuo and Ochman 2009). Ultimately, this dynamic has the potential to drive the irreversible loss of previously essential genes during prolonged colonization of a host and may enable host dependence to form more rapidly in prokaryotic organisms than in eukaryotes, which do not have a strong deletion bias (Denver *et al.* 2004; Kuo and Ochman 2009; Dyall *et al.* 2014). Consistent with this dynamic, host-restricted *Burkholderia* genomes evolving at lower N_e_ are indeed substantially smaller than free-living genomes (Mahenthiralingam *et al.* 2005; Carlier and Eberl 2012).

The lack of mutational bias towards AT bases observed in *B. cenocepacia* has not been seen previously in non-mutator MA lineages of any kind (Lind and Andersson 2008; Lynch *et al.* 2008; Denver *et al.* 2009; Keightley *et al.* 2009; Ossowski *et al.* 2010; Lee *et al.* 2012; Sung *et al.* 2012a; b). However, selection and/or biased gene conversion must still be invoked to explain the high %GC-content in *B. cenocepacia* (Hershberg and Petrov 2010; Hildebrand *et al.* 2010). Of these two explanations, selection favoring GC-content may be the more influential force, given that there is no evidence for increased %GC-content in recombinant genes of *Burkholderia*, despite its prevalence in other bacteria (Lassalle *et al.* 2015). It is also notable that similar substitution biases can be observed at polymorphic sites of several MLST loci shared across *B. cenocepacia* isolates (Jolley and Maiden 2010). Specifically, A:T>C:G transversions are more common than G:C>T:A transversions, and the rates of G:C>A:T and A:T>G:C transitions are nearly indistinguishable at six of the seven loci (see Figure S7). However, the evolutionary mechanism of these substitution biases are uncertain given the potential for ongoing recombination and/or natural selection to influence polymorphisms at these sites in conserved housekeeping genes (Lynch *et al.* 2008; Duret and Galtier 2009; Raghavan *et al.* 2012; Zhu *et al.* 2014).

In principle, a decreased rate of G:C>A:T transition mutation relative to other bacteria could be achieved by an increased abundance of uracil-DNA-glycosylases (UDGs), which remove uracils from DNA following cytosine deamination (Pearl 2000), or by a lack of cytosine methyltransferases, which methylate the C-5 carbon of cytosines and expose them to increased rates of cytosine deamination (Kahramanoglou *et al.* 2012). However, *B. cenocepacia* HI2424 does not appear to have an exceptionally high number of UDGs, and it does contain an obvious cytosine methyltransferase homolog, suggesting that active methylation of cytosines does occur in *B. cenocepacia*. Extending these methods to more genomes with high %GC-content will be required to determine whether a lack of AT-mutation bias is a common feature of GC-rich genomes.

Perhaps the most important finding from this study is that both mutation rates and spectra vary significantly among the three autonomously replicating chromosomes that make up the *B. cenocepacia* genome (Figure 3). Our data demonstrate that base-substitution mutation rates vary significantly among chromosomes, but not in the direction predicted by comparative studies on sequence divergence (Mira and Ochman 2002; Cooper *et al.* 2010; Lang and Murray 2011; Agier and Fischer 2012; Morrow and Cooper 2012). Specifically, we find that base-substitution mutation rates are highest on the primary chromosome (Figure 3A,B), where evolutionary rates are lowest. Thus, purifying selection must be substantially stronger on the primary chromosome to offset the effect of an elevated mutation rate.

The spectra of base-substitutions also differed significantly among chromosomes. Specifically, A:T>C:G transversions are more than twice as likely to occur on chr1 as elsewhere, and G:C>T:A transversions are more than twice as likely to occur on the chr3 (Figure 3C). One possible explanation for the increased rate of G:C>T:A transversions on chr3 is that they can arise through oxidative damage (Michaels *et al.* 1992; Lee *et al.* 2012) and may be elevated late in the cell cycle when intracellular levels of reactive oxygen species are high (Mira and Ochman 2002; Stamatoyannopoulos *et al.* 2009; Chen *et al.* 2010). Thus, because tertiary chromosomes are expected to be replicated late in the cell cycle (Rasmussen *et al.* 2007), we would expect these elevated rates of G:C>T:A transversions on chr3. Of course, if this explanation were accurate, we would also observe and increased rate of G:C>T:A transversions in late-replicated regions of chr1 and chr2. Although the low number of total G:C>T:A transversions observed in this study prevents us from statistically distinguishing G:C>T:A transversion rates between late and early replicated regions of chr1 and chr2, the rate of G:C>T:A transversions is higher in late replicated regions of chr1 and chr2 (see Figure 5A), a remarkable finding considering that early replicated genes on chr1 and chr2 are expressed more, which has been shown to induce G:C>T:A transversions independently of replication (Klapacz and Bhagwat 2002; Kim and JINKS-ROBERTSON 2012; Alexander *et al.* 2013). Thus, we suggest that late replicating DNA is inherently predisposed to increased rates of G:C>T:A transversions, possibly due to increased exposure to oxidative damage (Michaels *et al.* 1992), variation in nucleotide-pool composition (Kunkel 1992; Zhang and Mathews 1995), or variation in DNA-repair mechanisms (Hawk *et al.* 2005; Courcelle 2009).

A mechanism of an increased A:T>C:G transversion mutation rate on the primary chromosome is less clear, but a decreased rate of A:T>C:G transversions in a late replicating reporter relative to that on an intermediate replicating reporter has been demonstrated previously in *Salmonella enterica* (Hudson *et al.* 2002). Thus, it is possible that the rate of this form of transversion is increased in early replicating DNA, or that it is primarily caused by other forms of mutagenesis (Klapacz and Bhagwat 2002). A:T>C:G transversion rates in early replicating regions of chr1 and chr2 support the former hypothesis, as early replicated regions of chr1 and chr2 experience the highest rates of A:T>C:G transversions (see Figure 5B). The alternative mechanism of transcriptional mutagenesis seems less likely as A:T>C:G transversions occurred frequently in non-coding DNA relative to other substitution types (see Figure S8).

In summary, this study has demonstrated that the GC-rich genome of *B. cenocepacia* has a relatively low mutation rate, with a mutation spectrum that lacks an AT-bias and is biased toward deletion. Moreover, both the rate and types of base-substitution mutations that occur most frequently vary by chromosome, likely related to replication dynamics, the cell cycle, and transcription (Klapacz and Bhagwat 2002; Cooper *et al.* 2010; Merrikh *et al.* 2012). Although this study has broadened our understanding of mutation rates and spectra beyond that of model organisms, whether the observed mutational traits are common to all GC-rich genomes with multiple replicons, or are merely species-specific idiosyncrasies will require a more thorough investigation across a more diverse collection of GC-rich and multi-replicon bacterial genomes. Ultimately, by better understanding the core mutational processes that generate the variation on which evolution acts, we can aspire to develop true species-specific null-hypotheses for molecular evolution, and by extension, enable more accurate analyses of the role of all evolutionary forces in driving genome evolution.

## ACKNOWLEDGMENTS

We thank Kenny Flynn for helpful discussion and Brian VanDam for technical support. This work was supported by the Multidisciplinary University Research Initiative Award from the US Army Research Office (W911NF-09-1-0444 to ML, P. Foster, H. Tang, and S. Finkel); and the National Science Foundation Career Award (DEB-0845851 to VSC).

